# A programmable, selection-free CRISPR interference system in *Staphylococcus aureus*

**DOI:** 10.1101/2024.11.27.625599

**Authors:** Roni Miah, Mona Johannessen, Morten Kjos, Christian S. Lentz

**Affiliations:** Department of Medical Biology and Centre for New Antibacterial Strategies (CANS), UiT- The Arctic University of Norway, 9019, Tromsø, Norway; Faculty of Chemistry, Biotechnology and Food Science, Norwegian University of Life Sciences, Ås, Norway

## Abstract

Common dCas9-based CRISPR interference (CRISPRi) system for gene regulation requires antibiotic selection and exogenous inducer molecules, posing significant challenges when applied in *in vivo* bacterial infection models. Using *Staphylococcus aureus* as a model organism, we have developed a programmable, plasmid-based, but selection-free (ppsf)-CRISPRi system that is based on the pCM29- plasmid which is stable without antibiotic selection. In this ppsf-CRISPRi system, dCas9 expression is regulated by an endogenous virulence gene promoter, and sgRNA expression is driven by a constitutive promoter eliminating the need for exogenous inducer molecules. The system was programmed to silence the expression of genes encoding the virulence factor coagulase or peptidoglycan hydrolase autolysin, whenever their respective endogenous promoter was activated. The selection-free functionality was confirmed over at least 27 generations and verified by qPCR and phenotypic assays depending on the protein target, including coagulation of rabbit plasma and THP-1 macrophage cell infection *in vitro* as well as *in vivo* infection of *Galleria mellonella* larvae, in each case phenocopying the observations made using transposon mutant strains. The system is suitable for long-term studies of *S. aureus* pathogenesis *in vitr*o or *in vivo* and represents a blueprint for the development of similar ppsf-CRISPRi systems in other bacterial species.

## GRAPHICAL ABSTRACT

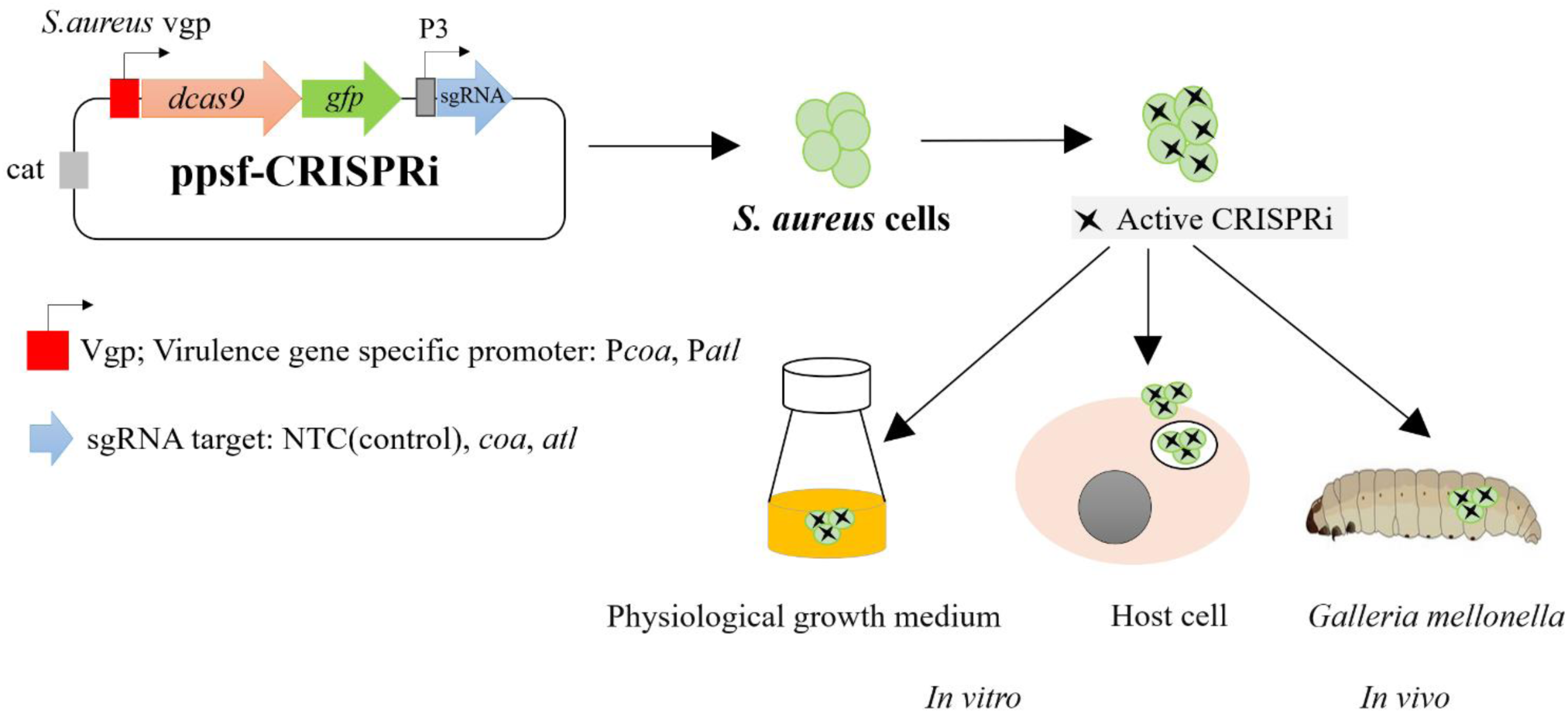

## INTRODUCTION

The unfolding antimicrobial resistance crisis urges the development of new antimicrobial treatment options for antibiotic-resistant priority pathogens, including methicillin-resistant *Staphylococcus aureus* (MRSA) (1)*. S. aureus* colonizes the nose in approx. 30% of the human population and can cause a broad range of both local and systemic, life-threatening systemic infections (2–5). Assigning gene function at the dynamic host-pathogen interface is critical for understanding the mechanisms of bacterial pathogenesis and evaluating potential drug targets. Traditional approaches in functional genomics of bacteria use insertion or deletion mutagenesis methods that are labour-intensive and can only be applied to non-essential genes. Recent studies have utilized CRISPR (Clustered Regularly Interspaced Short Palindromic Repeats) technologies, and particularly CRISPR interference (CRISPRi) have found wide applications in various bacterial species (6–15). CRISPRi gene-silencing utilizes an inactive Cas9 protein (dCas9) and customized single-guide RNA (sgRNA) complex to bind to a specific gene locus for effectively blocking transcription (6). The sgRNA spacer sequence of the CRISPRi tool can be easily modified to target the specific genomic regions of interest (16, 17) expanding this system’s utility in basic and applied microbial research. Furthermore, CRISPRi-seq uses pooled CRISPRi libraries and sequencing for genome-wide quantification of gene fitness that holds significant promise for studying bacterial pathogenesis (11–15).

The delivery of CRISPRi systems to bacteria has traditionally relied on replicative plasmids, requiring both antibiotic selection and exogenous inducers to achieve effective CRISPRi-based gene silencing (6–8, 12–15, 18, 19). However, antibiotics used for selection can cause cellular changes even at subinhibitory levels, and resistance mechanisms, such as antibiotic efflux or protein mutations, come with functional costs that can affect bacterial fitness (20–31). Furthermore, long-term *in vivo* experiments face challenges, such as the degradation of antibiotics over time, difficulties in antibiotic delivery (32). Antibiotic administration to maintain plasmid stability in animal studies is unreliable, as selective antibiotic levels may not reach all infection sites and could affect the animal’s microbiota, confounding experimental results (32). Additionally, some antibiotics do not effectively enter host cells, hindering plasmid maintenance in intracellular bacteria (33–36). The same pharmacokinetic limitations associated for antibiotics and CRISPR plasmid maintenance in experimental model systems of bacterial pathogenesis and infection also apply for exogenous inducer molecules, such as IPTG (isopropyl-β-D-thiogalactopyranoside) (37, 38, 13) or tetracyclines (39, 38), which are commonly used in the classical inducible CRISPRi (6, 13, 15, 18, 19, 40).

One approach to eliminate the need for antibiotic selection and inducer molecules is to integrate the complete, customized CRISPRi system into the bacterial chromosome - a method known as mobile-CRISPRi, which has been implemented in *Pseudomonas aeruginosa* (41, 42). CRISPRi system has also been optimized by integrating the inducible *dcas9* cassette into the genome of *S. aureus* (43, 44) and *S. pneumoniae* (40). However, chromosomal integration may induce other undesired phenotypes such as impaired growth (43) or other defects (45, 16, 17). Furthermore, construction of chromosomal integration strains is labor intensive in many species, including *S. aureus*, limiting the possibility of upscaling the cloning process (e.g., for construction of libraries for CRISPRi-seq).

Previously, we developed an inducer-free, endogenous virulence gene promoter-controlled, two plasmid-based CRISPRi system in methicillin-resistant *S. aureus* (46). However, as this system requires antibiotic selection, it is for that reason less suitable for *in vivo* studies. Here, we developed a programmable, plasmid-based, but selection-free (ppsf)-CRISPRi system in plasmid pCM29 that is both selection-free and exogenous inducer-independent. We used *S. aureus* as a model organism for demonstrating the proof-of-concept validation of the ppsf-CRISPRi system. In this study, the system leverages the promoters of the virulence-associated genes *coa* (encoding coagulase) and *atl* (encoding autolysin) to regulate dCas9 expression (47, 48). The retention of pCM29 plasmid without antibiotic selection enables gene silencing in the absence of external inducers and antibiotics. The system also includes a fluorescent reporter gene downstream of *dcas9* that indicates when dCas9 is expressed to activate the CRISPRi system. Proof-of-concept validation was achieved by silencing genes involved in various phenotypes and by demonstrating the system’s utility in the study of both *in vitro* and *in vivo* models of *S. aureus* infection.

## MATERIALS AND METHODS

### Bacterial strains, media, growth conditions, and plasmids transformation

*E. coli* and *S. aureus* USA 300 LAC or JE2 wild-type (WT), along with their genetically engineered strains, and the plasmids used in this study are listed in **Table S1** (bacterial strains) and **Table S2** (plasmids). The growth media used for *E. coli* were Lysogeny Broth (LB) (BD Difco™) or LB agar (BD Difco™), and for *S. aureus* were Tryptic Soy Broth (TSB) (BD Difco™) or Tryptic Soy Agar (TSA) (BD Difco™). RPMI-1640 was purchased from Sigma-Aldrich (R8758) and was supplemented with 10% FBS (Sigma-Aldrich) and is designated as RPMI+. Depending on the experimental conditions, *E. coli* cells were cultured in LB broth with agitation or on LB agar plates at 37 or 27°C. Ampicillin was used at a final concentration of 100 µg/ml in LB broth or agar plates for the selection of recombinant plasmid-transformed *E. coli* colonies. *S. aureus* was routinely grown at 37°C on TSA plates and in TSB or RPMI+ with shaking (220 rpm). Chloramphenicol (10 µg/ml) was used for the selection of recombinant pCM29-backbone plasmid-transformed *S. aureus* strains. Erythromycin (5 µg/ml) and chloramphenicol (10 µg/ml) were used for maintenance of pLOW and pVL2336 backbone plasmids, respectively, in *S. aureus* strains. IPTG was used at 250 µM (final concentration) for *lac*- promoter-induced dCas9 expression in the classical CRISPRi system. Chemically competent *E. coli* IM08B cells were prepared according to Chang Y et al., 2017 (49) and routinely used for transformation of constructed plasmid according to standard heat shock protocol (50). *S. aureus* USA 300 LAC was transformed with plasmid DNA isolated from IM08B (51) by electroporation.

Preparation of *S. aureus* electrocompetent cells and electroporation were performed as described before (52).

### Generation of ppsf-CRISPRi constructs

To generate ppsf-CRISPRi constructs, the *dcas9* gene was amplified from pLOW-P*lac*-*dcas9* (18) using the primer sets RM 27/28 (**Table S3**). The amplified fragment was digested with Bsu36I (New England Biolabs) and ligated into the corresponding site of a previously constructed pCM29-based sgRNA expression vector (46) using T4 Ligase (New England Biolabs). The ligation mixture was then transformed into *E. coli* IM08B cells, plated on LB agar containing 100 µg/ml ampicillin, and incubated overnight at either 27°C or 37°C. The correct construct was confirmed through restriction mapping, PCR, and whole plasmid sequencing. This resulted in the ppsf-CRISPRi constructs with *pbp1*, *coa*, *atl* and non-targeting control (NTC) as the sgRNA (**Table S2**). In these constructs, the expression of *dcas9* and *gfp* is regulated by the virulence gene promoters of coagulase (*coa*) and autolysin (*atl*), while sgRNA expression is driven by a constitutive P3 promoter. The sequences for the sgRNA base-pairing regions are listed in **Table S4**.

### Bacterial growth and GFP fluorescence analysis

Bacterial growth (OD600) and GFP fluorescence were measured in a microplate assay on a Synergy H1 Hybrid Reader (BioTek). *S. aureus* strains were grown overnight with shaking at 37 °C, and *E. coli* strains were grown at 37 or 27 °C as specified. Cell cultures were diluted to an OD600 of 0.1 in their respective fresh medium (TSB or RPMI+ for *S. aureus* and LB for *E. coli* strains, respectively). Two μL of these diluted overnight cultures were inoculated into 298 μL of each respective fresh medium in a 96-well flat-bottom polystyrene tissue culture plate (Falcon). The plate was incubated directly in a Synergy H1 Hybrid Reader (BioTek) microtiter plate reader at 37 or 27°C as specified with constant double orbital shaking (400 rpm for 5 s) in-between measurements. Optical density (OD600) and GFP fluorescence (excitation 479, emission 520) were measured at 1-hour intervals for up to 24 hours. GFP fluorescence was recorded as relative fluorescence units (RFU, i.e. relative to the internal standard in the instrument). The OD600 and GFP fluorescence values from 3 biological replicates that were each derived from 3 technical replicates were averaged and corrected from blank wells containing only medium for each strain.

### Plasmid copy number determination by quantitative PCR (qPCR)

*S. aureus* ppsf-CRISPRi strain, MR36 and the previously constructed fluorescence reporter strain MR11 (**Table S1**) were initially grown in TSB medium with chloramphenicol (CHL). Cultured cells were then subjected to a three-step serial culture process in TSB with 24-hour intervals (denoted passages 1,2 and 3) in the presence or absence of CHL. From each passage, samples were harvested and subjected to total DNA isolation using the ZymoBIOMICS DNA miniprep kit (Zymo Research) following the manufacturer’s instructions and subjected to real-time PCR assay performed with PowerTrack SYBR Green Master Mix (Applied Biosystems™) containing 0.5 μM each primer set. After cycling, melt curves analysis was performed between 70°C and 90°C. All quantitative PCR (qPCR) data were analyzed using LightCycler® 96 system software version 1.1 (Roche Diagnostics). The plasmid copy number per cell was defined as the ratio of the plasmid to chromosomal DNA copies and was calculated using the formula 2^-ΔCq^, where -ΔCq represents the difference in quantification cycles between the plasmid gene (*gfp*) and the chromosomal gene (*groEL*) (53). The qPCR primers are listed in **Table S3**. qPCR was performed in three technical replicates from their three biological replicates.

### RNA purification, reverse transcription, quantitative PCR (qPCR), and coagulase test

*S. aureus* ppsf-CRISPRi (NTC) strain MR35 and the ppsf-CRISPRi (*coa*) strain MR36 (**Table S1**) were grown in TSB medium with CHL. Cultured cells were then subjected to a three-step serial culture process in TSB and RPMI+ medium with 24-hour intervals (denoted passages 1,2 and 3) in the presence or absence of CHL. Exponential cell cultures (OD600 of 1) from passage 3, were harvested and subjected to total mRNA isolation. Harvested cells were diluted in PBS (D8537, Sigma) and adjusted to McFarland 0.5 (i.e about 10^8^ cells/ml). Subsequently, cells were treated with 2× volume of RNAprotect Bacteria Reagent (QIAGEN) for 10 min at room temperature and collected as pellets by centrifugating for 10 min at 5,000×g. Lysostaphin, 1 µg/µL, and lysozyme, 10 µg/µL were added as final concentration to the pellets and subsequently prepared suspension was incubated at 37 °C for 30 minutes for lysis of bacterial cells before RNA isolation. RNA extraction was performed using a RNeasy Mini Kit (QIAGEN). The remaining DNA in the isolated RNA was degraded by DNase I (Sigma-Aldrich). DNA-free RNA was subjected to reverse transcription (RT) with the High-Capacity cDNA Reverse Transcription (RT) Kit (Applied Biosystems™). RNA extraction, DNase treatment, and RT were performed according to the manufacturer’s direction. Synthesized cDNA solution was 10,000-fold diluted with RNase-free water, and 2μl of the diluted cDNA solution was subjected to real-time PCR assay performed with PowerTrack SYBR Green Master Mix (Applied Biosystems™) containing 0.5 μM each primer set. After cycling, melt curves analysis was performed between 70°C and 90°C. All quantitative PCR (qPCR) data were analyzed using LightCycler® 96 system software version 1.1(Roche Diagnostics). Relative coagulase expression was calculated according to the 2^−ΔΔCt^ method after normalization by 16SrRNA (rrsA). The qPCR primers are listed in **Table S3**. qPCR was performed in three technical replicates from their three biological replicates.

For the coagulase test, exponential cell cultures (OD600 of 1) from passage 3 were diluted in PBS and adjusted to McFarland 0.5. Subsequently, 0.5 ml of each McFarland-adjusted culture was added to 0.5 ml of rabbit plasma (Bio-Rad). The samples were incubated in a water bath at 37°C for 4 h. The level of coagulation was verified by tipping the tubes to a 45° angle. A negative control (NC) sample contained medium only. A test was considered positive if the plasma in the tube formed a coherent clot. All experiments were repeated three times as independent biological replicates to examine the reproducibility.

### THP-1 cell culture

The human macrophage cell line THP-1 cells (TIB-202, ATCC) were cultured in RPMI+ media supplemented with 1% penicillin/streptomycin (Gibco™). Before the infection assay, THP-1 monocyte differentiation was initiated by incubating the cells with 10 ng/mL phorbol 12-myristate 13- acetate (Sigma-Aldrich) for 2 days at 37°C in a 5% CO2 environment. Following this 2-day differentiation period, the culture medium was aspirated, and the cells were washed three times with PBS. Subsequently, the cells were further incubated for 1 day in a fresh RPMI+ medium (supplemented with 10 µg/mL CHL when appropriate).

### Time-course THP-1 infection and cytotoxicity assay

For THP-1 infection and cytotoxicity assay, *S. aureus* ppsf-CRISPRi (NTC) strain MR 35 and the ppsf- CRISPRi (*atl*) strain, MR37 (**Table S1**) were grown in TSB medium with CHL. Cultured cells were then subjected to a three-step serial culture process in TSB medium with 24-hour intervals (denoted passages 1,2 and 3), either in the presence or absence of CHL. From passage 3, exponential cell cultures (OD600 of 1) were subjected to THP-1 infection assay. Alongside, overnight cultures of the *S. aureus* USA 300_LAC or JE2 (WT) and the autolysin transposon mutant strain, JE2 (*atl*::Tn) in TSB were sub-cultured into fresh TSB and grown to an OD600 of 1. Prepared *S. aureus* cells were diluted in PBS and adjusted to McFarland 0.5 and added to each well of seeded THP-1 macrophage cells at a multiplicity of infection (MOI) of 10. The host cells were infected for 1 hour, washed with PBS, and incubated in RPMI+ supplemented with 50µg/mL gentamicin to kill extracellular bacteria, and 10µg/mL CHL was added when appropriate.

For the time-course infection assay, at designated time points, the media were discarded, the mammalian cells were washed with PBS and lysed with 0.01% Triton X-100 in PBS, and the lysates were serially diluted and plated on TSA plates (supplemented with 10µg/mL CHL when appropriate). After 24 hours of incubation at 37°C, the bacterial colonies were counted.

For the time-course THP-1 cell cytotoxicity assay, at designated time points, the media were discarded, the cells were washed with PBS and MTT (3-(4,5-dimethylthiazol-2-yl)-2,5-diphenyltetrazolium bromide reagent, Invitrogen) was diluted (0.5mg/ml) in the RPMI+ cell culture media and added to the mammalian cells, followed by incubation at 37°C for 2 hours. The wells were washed twice with PBS, and dimethyl sulfoxide (DMSO) was added to solubilize the formazan crystals. Alongside these samples, control assays with uninfected cells (in the presence and absence of CHL) were conducted to ensure assay validity. The absorbance at 520 nm was measured using a BioTek Synergy H1 plate reader.

### Galleria mellonella infection assay

*Galleria mellonella* larvae were obtained from Reptilutstyr AS (Norway). For infection experiments, *S. aureus* ppsf-CRISPRi (NTC) strain, MR39 and ppsf-CRISPRi (*atl*) strain, MR40 (**Table S1**) were grown in TSB medium with CHL. The bacterial cells were then subjected to a three-step serial culture process in TSB medium with 24-hour intervals (denoted passages 1,2 and 3) in the absence of CHL. From passage 3, exponential cell cultures (OD600 of 1) were subjected to *Galleria* infection assay. Alongside, overnight cultures of the *S. aureus* USA 300_ LAC or JE2 (WT) and the autolysin transposon mutant strain, JE2 (*atl*::Tn) in TSB were sub-cultured into fresh TSB and grown to an OD600 of 1. Cells were washed twice in PBS, suspended in PBS, adjusted to McFarland 0.5, and diluted in PBS to a final concentration of 10^5^ cells/ml. Larvae of approximately equal weight were inoculated with 10 µL of this bacterial suspension, resulting in an infection dose of about 10^3^ cells/larva. Bacteria were injected into the hemocoel of the larvae between the last pair of legs using a 30-G syringe microapplicator (0.30 mm [30 G] × 8 mm, BD Micro-Fine Demi). As a control, larvae were mock- inoculated with 10 µL PBS. The larvae were then placed in 9.2 cm Petri dishes, and incubated at 37°C in darkness, and survival of the larvae was observed every 8 hours at 37°C. Larvae were regarded dead when they were not moving upon repeated physical stimulation. Each experiment was independently replicated at least three times using 3 biological replicates of each strain.

## RESULTS

### Construction of the ppsf-CRISPRi system in *S. aureus*

Antibiotic selection is essential to maintain the activity of our IPTG-inducible two plasmid-based CRISPRi system in *S. aureus*, as demonstrated by the reduced ability to induce growth inhibition phenotype through silencing of *pbp1* in the absence of selection pressure (**Figure S1**). Inspired by previous reports on the stability of pCM29 without chloramphenicol selection in *S. epidermidis* and *S. aureus* SH1000 (54), we wanted to construct a ppsf-CRISPRi system in *S. aureus* using pCM29. Attempts to clone ppsf-CRISPRi (*pbp1*) in *E. coli* IM08B at 37°C resulted in a truncation of the *dcas9* gene from 4107 to 559 bp, as confirmed by whole plasmid sequencing. This modification, likely due to dCas9-related toxicity as previously reported (46), rendered the constructs non-functional in *S. aureus* (**Figure S2**). However, the construct could be successfully created when the cloning was performed at 27°C. At this lower temperature, expression of the *S. aureus* gene promoters in *E. coli* was drastically reduced and there was no evidence of dCas9-based toxicity (**Figure S3**). Thus, we were able to construct stable ppsf-CRISPRi plasmids in *E. coli* IM08B (**Figure S4**, **Table S2**) that were directly transformed into *S. aureus* USA300 LAC strains for functional analysis.

### Evaluation of pCM29-plasmid backbone retention in *S. aureus* without antibiotic selection

To evaluate whether the ppsf-CRISPRi constructs are stable without antibiotic selection, we first used the plasmid-encoded fluorescence reporter as a proxy for plasmid stability and monitored fluorescence over 24h of growth in TSB and RPMI+, a common mammalian cell-culture medium, in the presence and absence of chloramphenicol. GFP fluorescence in ppsf-CRISPRi strain MR36, as well as in fluorescence reporter strain MR11 (**Table S1**) remained stable in the absence of chloramphenicol in both media (**Figure 1A-B**). To confirm the long-term presence of the plasmid, we determined the plasmid copy number across three serial culture passages with 24-hour intervals (equivalent to 27 generation times), both with and without chloramphenicol in TSB by qPCR (**Figure 1C**). The results indicate that both MR11 and MR36 strains maintained a consistent number of plasmids per cell, regardless of the presence or absence of chloramphenicol over the passage of culture days (**Figure 1D-E**). In conclusion, these results suggest the long-term stability of pCM29-plasmid backbone in *S. aureus* without antibiotic selection for at least 27 generations.

**Figure 1.**
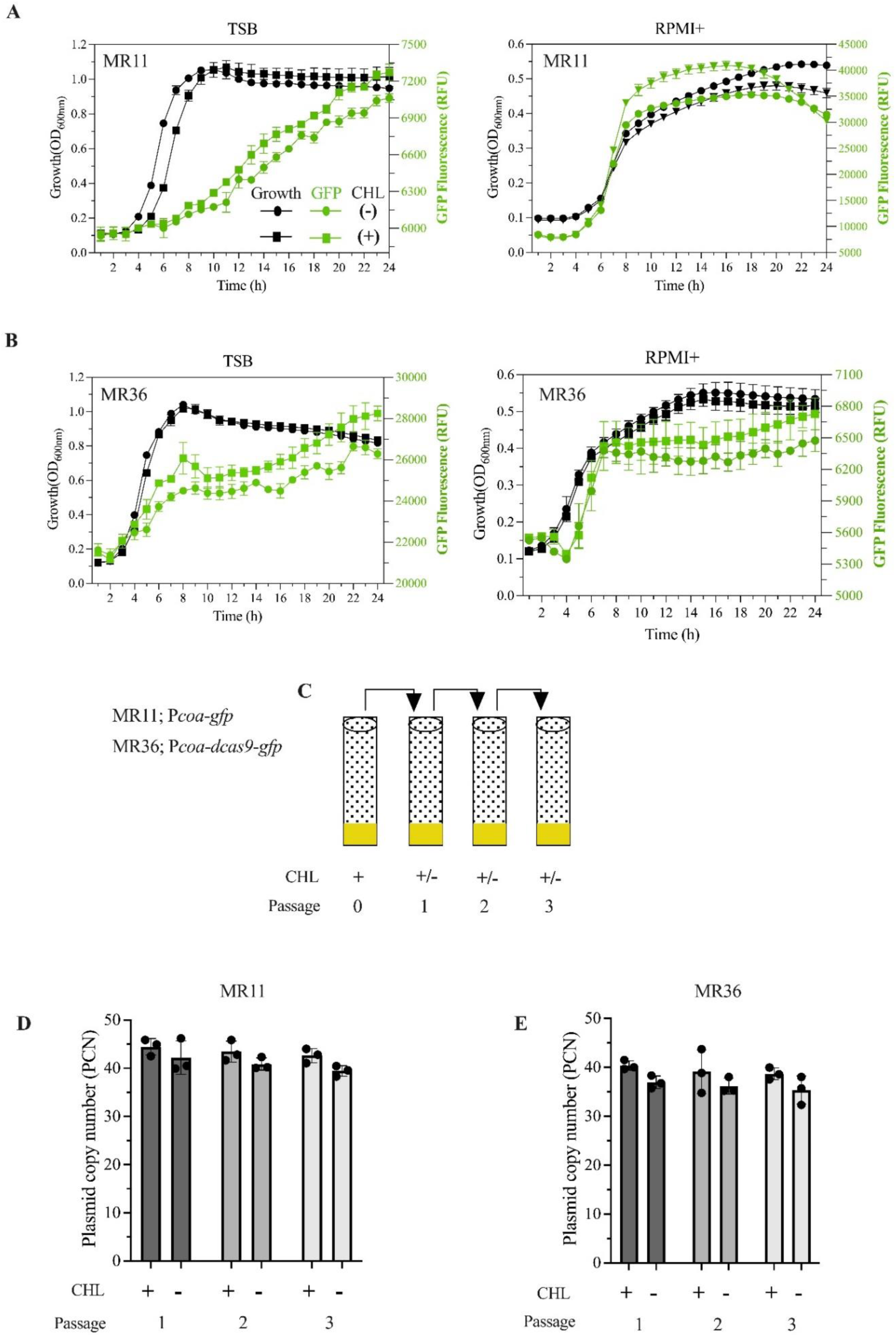
Retention of pCM29 plasmid in *S. aureus* USA 300 LAC without antibiotic selection. (A, B) Overlays of growth curves (OD600) and GFP fluorescence levels (in RFU) of *S. aureus* fluorescence reporter strains, MR11 (A, carrying pCM29-P*coa*-*gfp*) and ppsf-CRISPRi strain, MR36 (B, carrying pCM29-P*coa*-*dcas9*-*gfp*). Cultures were grown either in TSB or cell culture medium RPMI+ with (+) and without (-) chloramphenicol (CHL). (C) Schematic representation of serial culture passage with 24-hour intervals for the MR11 and MR36 strains. Cultures were initially grown in TSB medium with CHL and then passaged serially in TSB medium with (+) and without (-) CHL. (D, E) The copy numbers of pCM29-P*coa*-*gfp* in MR11 (D) and pCM29-P*coa*-CRISPRi(*coa*) in MR36 (E) per cell were determined using qPCR across three serial passages of cells cultured in the presence or absence of CHL. Data show means ± SD of n = 3 biological replicates (each recorded with three technical replicates).

### ppsf-CRISPRi system allows specific and robust interference with *coa* gene expression and coagulase function

We next aimed to evaluate whether the ppsf-CRISPRi system could effectively target specific gene expression and related cellular phenotypes even in the absence of antibiotic selection. As a model target, we chose coagulase, which is a critical virulence factor responsible for converting fibrinogen to fibrin, thereby causing plasma clotting (55,48). To assess the repression of coagulase transcription and function, we used ppsf-CRISPRi strain MR36 with *coa* as the sgRNA target and MR35 as a non-targeting control (NTC) (**Table S1**). In both strains, dCas9 expression was regulated by the P*coa* promoter. Both MR35 and MR36 were subjected to three consecutive culture passages at 24-hour intervals in TSB and RPMI+ media, with and without chloramphenicol (**Figure 1C**) prior to assessing expression of *coa* using qPCR.

The results showed that *coa*-targeting MR36 strain exhibited a significant reduction in *coa* expression compared to the control strain MR35, regardless of the growth medium and presence or absence of chloramphenicol (**Figure 2A**). The sustained silencing of the *coa* gene in strain MR36 (even after three 24-h passages in the absence of selection) led to a functional deficiency in coagulation of rabbit plasma (**Figure 2B**), suggesting that coagulase activity was efficiently abrogated. In contrast, clotting occurred in both CRISPRi control strain (MR35) and WT strains. This demonstrates that the ppsf-CRISPRi system can robustly silence virulence genes like *coa*, effectively interfering with their biological function in *S. aureus* without the need for antibiotic selection.

**Figure 2.**
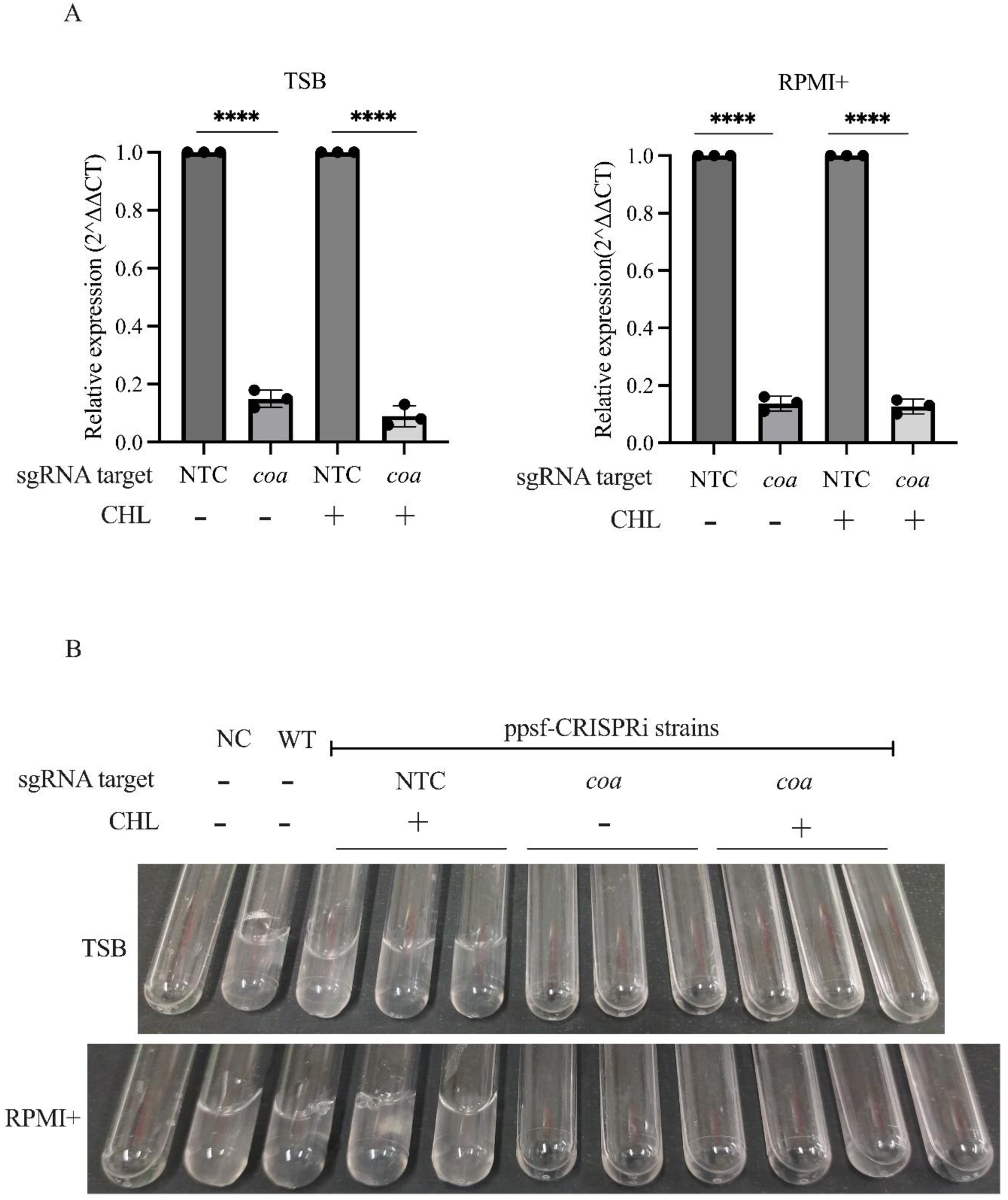
Silencing of *coa* expression and plasma coagulation using the ppsf-CRISPRi system. (A) The repression of *coa* mRNA of the indicated ppsf-CRISPRi strains third passage grown in TSB or RPMI+ with (+) and without (-) chloramphenicol (CHL). The sgRNA target used was *coa*. A non- targeting sgRNA was included as control (NTC). dCas9 expression was regulated by the P*coa* promoter. qPCR was performed on exponential cell cultures (OD600 of 1) harvested after passage three. The relative coagulase expression was calculated after normalization by 16S rRNA (*rrsA*). The data represents the mean ± SD of n = 3 biological replicates, each recorded with two technical replicates. Significance was tested for each construct against its NTC control pair by unpaired, two-tailed Student’s t-test. ****P < 0.0001. (B) The figure shows photographs of coagulase test tubes 4 hours after adding indicated third passage ppsf-CRISPRi strains to rabbit plasma. The coherent clot formation is indicative of coagulation (arrowheads). Three tubes for each strain are representative of three biological replicates. The negative control (NC) contained medium (TSB or RPMI+) with rabbit plasma, and WT represents the *S. aureus* USA 300 LAC strain with rabbit plasma.

### ppsf-CRISPRi system’s functionality in studying *S. aureus* infection of human cell line

To evaluate the utility of ppsf-CRISPRi system for studying *S. aureus* infection in human cell lines, we monitored the intracellular survival of bacteria in the human THP1 macrophage cell line, as well as THP1 cell viability over 48 h. As a model target, we chose to interfere with the *atl* gene, which encodes autolysin (Atl), a peptidoglycan hydrolase critical for cell wall degradation and division (56). The strains used included WT *S. aureus* USA300 LAC and two ppsf-CRISPRi constructs: one targeting *atl* (MR37) and a non-targeting control (NTC, MR35), both with dCas9 expression driven by the P*coa* promoter (**Table S1**). Additionally, WT *S. aureus* USA300_JE2 and its *atl* transposon mutant strain were used as controls. The ppsf-CRISPRi strains were subjected to the same 3-passage pre-cultivation in the presence/absence of antibiotic selection described earlier (**Figure 1C**) and THP-1 cell infection experiments were carried out in the presence or absence of chloramphenicol. Chloramphenicol itself had no effect on the viability of THP1 cells (**Figure S5**). The viability of macrophages infected with WT or non-targeting ppsf-CRISPRi control strain MR35 decreased markedly over the time, whereas macrophages infected with *atl*-deficient strains (both transposon mutant and ppsf-CRISPRi knockdown) had significantly higher viability (**Figure 3 A-C**, **left panel**). We observed that the levels of viable intracellular bacteria (CFUs) were reduced approx. 10-fold in the *atl* transposon mutant compared to the WT throughout the assay period (**Figure 3A**, **right panel**). Similarly, regardless of the presence or absence of chloramphenicol in the pre-cultivation period and throughout the experiment, the *atl*-targeting ppsf-CRISPRi strain MR37 exhibited a similar 1-log reduction of intracellular CFU when compared to WT and NTC strains (**Figure 3 B-C**, **right panel**).

**Figure 3.**
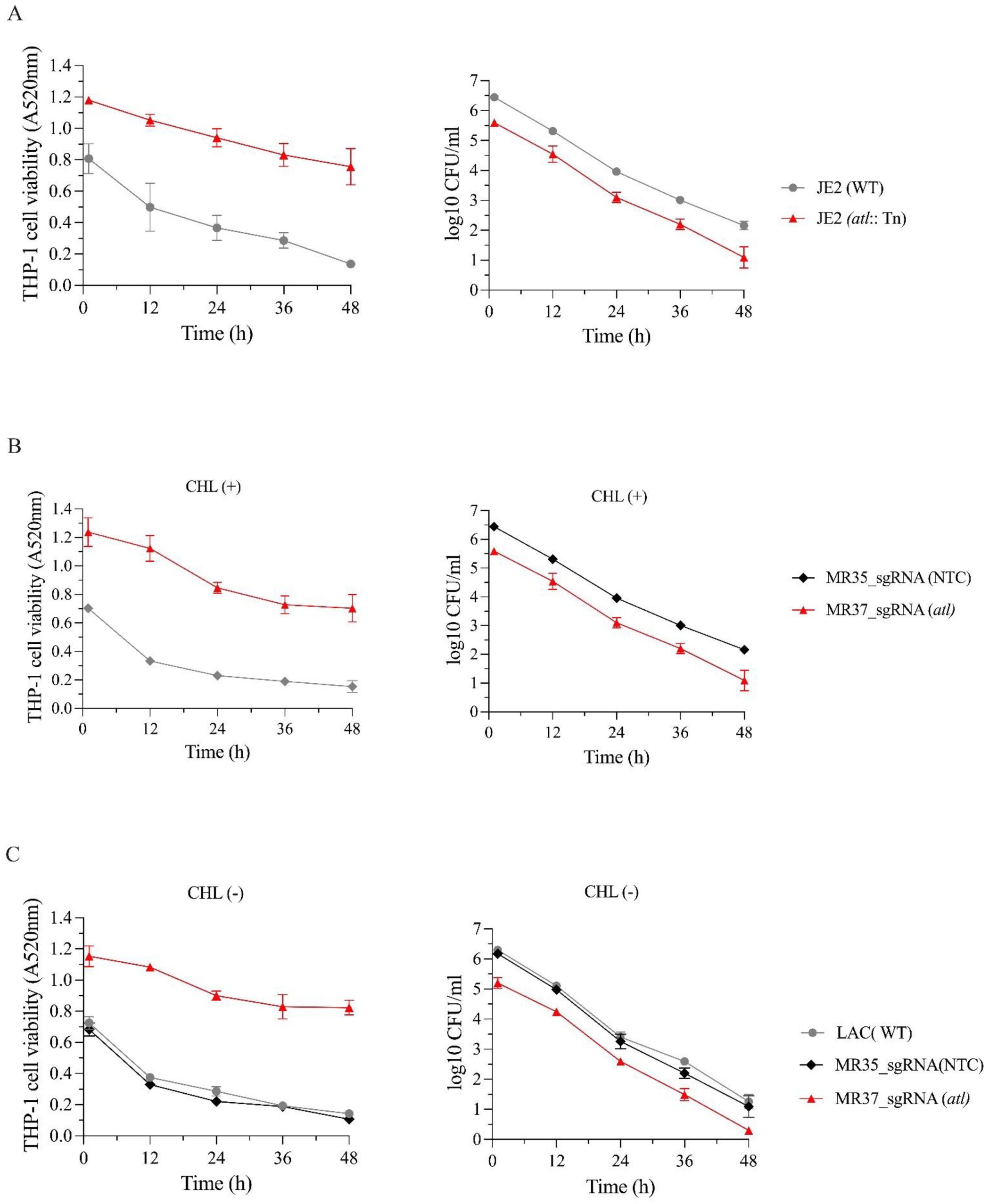
ppsf-CRISPRi system to study THP-1 macrophage infection with *S. aureus*. (A, B, and C) Time-course THP-1 cell viability (MTT) (left panel) and infection assay (right panel) following infection with the indicated *S. aureus_*JE2 or LAC WT, transposon mutant JE2 (*atl*::Tn), and ppsf-CRISPRi control strain (MR35, NTC) and strains targeting *atl* (MR37). For ppsf-CRISPRi constructs, the sgRNA target was *atl* or an NTC derived from the luciferase gene, dCas9 expression controlled by the P*coa* promoter. Before the infection assay, the indicated ppsf-CRISPRi strains were grown in TSB medium with CHL. They were then serially passaged in RPMI+ medium with (+) and without (-) CHL at 24-hour intervals for up to three culture passages. Following the final culture passage, the (+) CHL and (-) CHL cultivated strains were subjected to the time-course THP-1 infection in the presence and absence of CHL respectively. The data represents the mean ± standard deviation of three biological replicates.

Since the difference in intracellular CFU between *atl*-deficient and control strains manifests already at 1 h and the kinetics of the decline in CFU over time is similar across all strains, our data suggest that these outcomes are not due to reduced intracellular viability or faster killing of *atl*-deficient bacteria, but rather reflect reduced internalization. This finding aligns with previous research demonstrating that *S. aureus* autolysin mutant strains exhibit decreased binding to host cell ligands and reduced internalization into host cells (57, 47). Collectively, these results support the effectiveness of the ppsf- CRISPRi system in studying *S. aureus* infections in host cell infection or co-culture experiments without selection.

### The ppsf-CRISPRi system is functional during *Galleria mellonella* infection

To evaluate the utility of ppsf-CRISPRi system for studying *S. aureus* virulence in an *in vivo* setting, we used the *G. mellonella* infection model. *G. mellonella* larvae were infected with a P*atl*-driven ppsf- CRISPRi strain targeting *atl* (MR40) and a non-targeting control strain (MR39) (**Table S1**). Wild-type USA300_JE2 or LAC strains and corresponding *atl* transposon mutant were included as controls. Larval survival was monitored over 96 hours. Infections with either the WT strain or the non-targeting ppsf-CRISPRi control (MR39) resulted in 100% mortality within 96 hours (**Figure 4**). In contrast, larvae infected with the *atl* transposon mutant or the ppsf-CRISPRi *atl* knockdown strain (MR40) had a significantly higher survival rate. Approximately 30% of the larvae infected with these strains were still alive after 96 hours (**Figure 4**). This result show that disrupting *atl* gene function, either by transposon mutation or by CRISPRi knockdown, reduces the virulence of *S. aureus*, leading to increased larval survival. These data also demonstrate that the ppsf-CRISPRi system is indeed efficient in studying *S. aureus* infection in *G. mellonella* infection model without relying on antibiotic selection.

**Figure 4.**
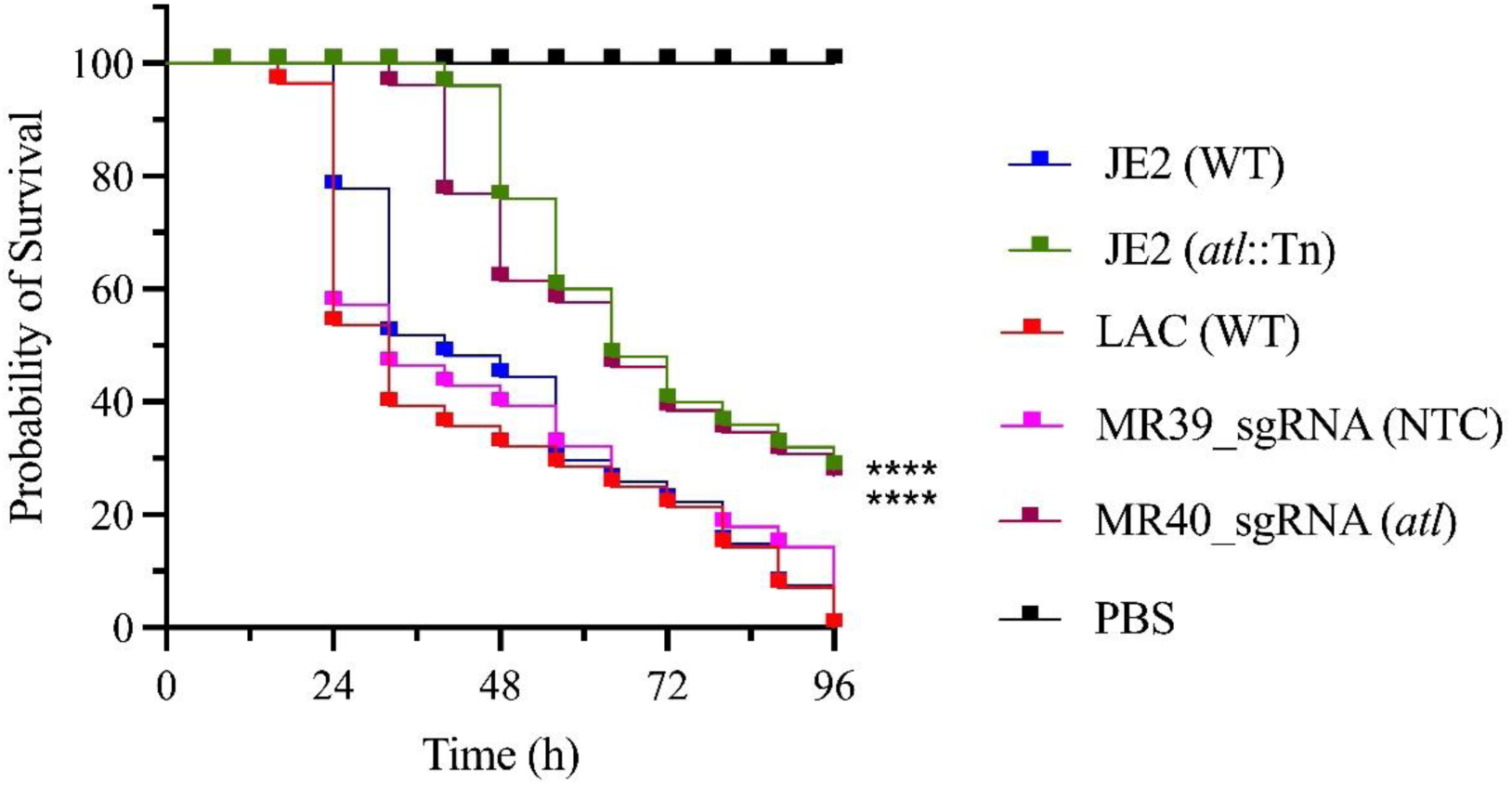
ppsf-CRISPRi system for studying *Galleria mellonella* infection with *S. aureus.* Kaplan– Meier (KM) survival plots of *G. mellonella* larvae after inoculation with indicated *S. aureus* WT, transposon mutant, and ppsf-CRISPRi strains. For ppsf-CRISPRi strains, dCas9 expression was controlled under autolysin gene promoter (P*atl*). Before the *G. mellonella* infection study, the ppsf- CRISPRi strains were cultured in TSB medium with CHL, and then passaged up to three times in antibiotic-free TSB medium at 24-hour intervals. The plots indicate the average of n=3 independent biological replicates (i.e. bacterial cultures originating from 3 different colonies), that were each used to infect 10 larvae per group. Mortality was monitored every 8 hours for 96 hours (N=180). Larvae injected with PBS served as a negative control. The asterisks in the graph indicate a significant difference in the survival of autolysin (*atl*) transposon mutant strain compared to their wild-type (WT) strain and *atl* silenced CRISPRi strain compared to their non-target CRISPRi control (NTC) or wild-type (WT) strain (**** p < 0.0001).

## DISCUSSION

CRISPRi-based approaches hold promise for the identification of virulence factors critical for bacterial pathogenesis and understanding the bottlenecks that pathogens encounter during infection (13). However, the widely used plasmid-based inducible CRISPRi system requires antibiotic selection for plasmid maintenance and inducer molecules for dCas9 expression. While functional, this classical system carried drawbacks for *in vivo* studies of bacterial pathogenesis due to its reliance on inducers and antibiotics, which can impact bacterial fitness and experimental outcomes. We explored an alternative approach for inducer independent and selection free CRISPRi system development. Previously, we developed an endogenous virulence gene promoter (vgp)-controlled CRISPR interference (vgp-CRISPRi) system in *S. aureus* (46). This system did not require external inducers, but still relied on antibiotic selection, similar to the classical inducible CRISPRi system. Despite the progress made with this vgp-CRISPRi system, the need for antibiotics introduced potential challenges for applying this system for *in vivo* studies of *S. aureus*. Inspired by previous reports on the stability of pCM29 plasmid without chloramphenicol selection in *S. epidermidis* and *S. aureus* SH1000 (54), we have developed the pCM29-based ppsf-CRISPRi system in the MRSA *S. aureus* strain, which was retained and active in *S. aureus* even in the absence of antibiotic selection for at least 27 generation times.

How can the ppsf-CRISPRi system be programmed? In the current study, the constructs were benchmarked against and designed to mimic knock-out strains. By placing dCas9 expression under control of the promoter that also control the target of the sgRNA (here *coa* or *atl*), our constructs were programmed to interfere with the expression of these virulence genes, whenever their promoter is activated. Of note, in all assay systems, the performance of the ppsf-CRISPRi system phenocopied the effects of the transposon mutants, demonstrating that it is a viable alternative to studies using knock- out strains.

An important practical consideration for future applications is the feasibility of strain construction. During the generation of this ppsf-CRISPRi construct, we encountered dCas9 toxicity-dependent plasmid modification issues in *E. coli* due to the expression of *dcas9* from the *S. aureus* gene promoter at 37°C. The molecular mechanism behind this dCas9 toxicity is unknown but may be related to the previously reported “bad-seed” sequence (58). Rostain et al. 2023 (58) demonstrated that when dCas9 is expressed along with guide RNAs carrying bad-seed sequences, it could bind to hundreds of off-target positions in *E. coli* bacterial genomes leading to silencing and toxicity. Plasmid modification caused by dCas9 toxicity in the cloning host *E. coli* remains a major challenge in designing customized CRISPRi circuits, as we have observed. Our findings underscore an important consideration for developing single-plasmid CRISPRi systems controlled by endogenous gene promoters in any target strains when *E. coli* is used as an intermediate cloning host. It is critical to verify whether the endogenous promoter driving dCas9 expression is active in *E. coli*. If the promoter is indeed active, strategies to reduce its activity and minimize dCas9 expression are essential to prevent dCas9 expression at toxic levels. By lowering the cloning temperature, we succeeded in mitigating that dCas9 toxicity-dependent plasmid modification issues and generated stable ppsf-CRISPRi constructs in *S. aureus*. The application of ppsf-CRISPRi system was successfully validated for both *in vitro* and *in vivo* studies of *S. aureus* without selection.

In addition to gene silencing, the ability to monitor promoter activity using a fluorescent reporter downstream of the *dcas9* gene provides a dual-function of ppsf-CRISPRi tool for studying both gene regulation and promoter dynamics under different environmental conditions. In theory, if gene promoters that control *dcas9* are only activated under specific conditions e.g. during infection, activation of CRISPRi system would then also be restricted to those conditions. Similarly, cell-to-cell heterogeneity in gene expression and promoter activity could be exploited for specific targeting of bacterial subpopulations.

One of the advantages of CRISPRi is that it allows the inducible knock-down of essential genes (18, 40, 43). With the ppsf-CRISPRi system, this would be feasible if the endogenous promoter is replaced again by an inducible promoter. Such a system could be operated free of antibiotic selection but would again rely on external inducer molecules. We anticipate that in the future the exploration of a variety of different exogenous gene promoters will allow ppsf-CRISPRi system to be programmed to silence essential genes (16) of several bacterial species. We envision that employing a weak constitutive promoter for dCas9 expression (41, 42) in broad-host-range, antibiotic-selection-free plasmids (59) will facilitate ppsf-CRISPRi approach across diverse bacterial species without the complications of chromosomal integration. The incorporation of Golden Gate cloning sites into the ppsf-CRISPRi construct will enhance the flexibility and efficiency of sgRNA cloning, facilitating the construction of desired CRISPRi strains in just a single cloning step. Of note, the system may further be programmed by also placing the sgRNA under control of promoter that control dCas9 expression.

In conclusion, the ppsf-CRISPRi system a versatile and efficient tool for exploring gene expression and regulation in *S. aureus* without the limitations associated with chromosomal integration, inducers, and antibiotic selection. We believe that the adaptability and flexibility of the ppsf-CRISPRi approach will enable precise, programmable gene silencing within specific bacterial species in a complex microbiome offering new insights into the functional dynamics of bacterial interactions at the host interface.

## AUTHOR CONTRIBUTIONS

Experiment design: All authors, Data acquisition: R.M., Data analysis: All authors, Original manuscript draft: R.M., Manuscript writing: All authors. Concept: R.M. and C.S.L. Funding: C.S.L.

## ACKNOWLEDGMENTS

We would like to thank Simen Hermansen, NMBU, for structure analysis of mutated dCas9 variants. We would also like to express our gratitude to Bhupender Singh and Julia Maria Kloos, UiT for their assistance in coordinating with Eurofins Genomics for the whole plasmid sequencing and Kjersti Julin, UiT for assistance with human cell culture.

---

**Figure S1.**
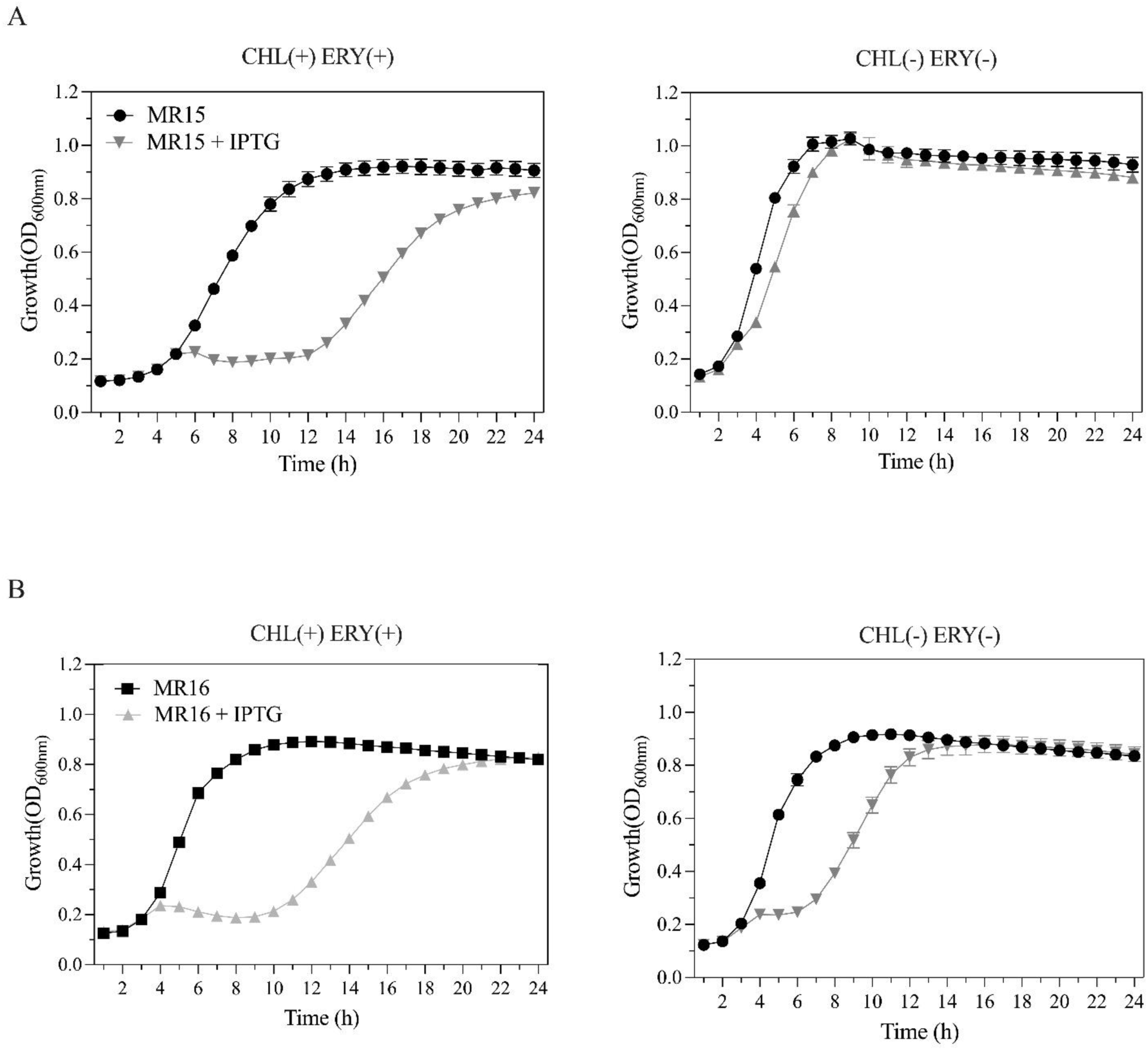
Antibiotic selection is essential for effectiveness of the classical CRISPRi system. Growth (OD600) of IPTG-inducible CRISPRi strains targeting the essential peptidoglycan biosynthesis gene *pbp1*. The expression of dCas9 was regulated through IPTG induction via *lac* promoter on plasmid pLOW, while sgRNA is constitutively expressed from either pVL2336 (MR15, A) or pCM29 (MR16, B). Cells were grown in TSB with (+) and without (-) antibiotic selection in the presence or absence of IPTG (250 µM). Data show mean ± SD of n = 3 biological replicates (each recorded with three technical replicates). CHL; chloramphenicol, ERY; erythromycin.

**Figure S2.**
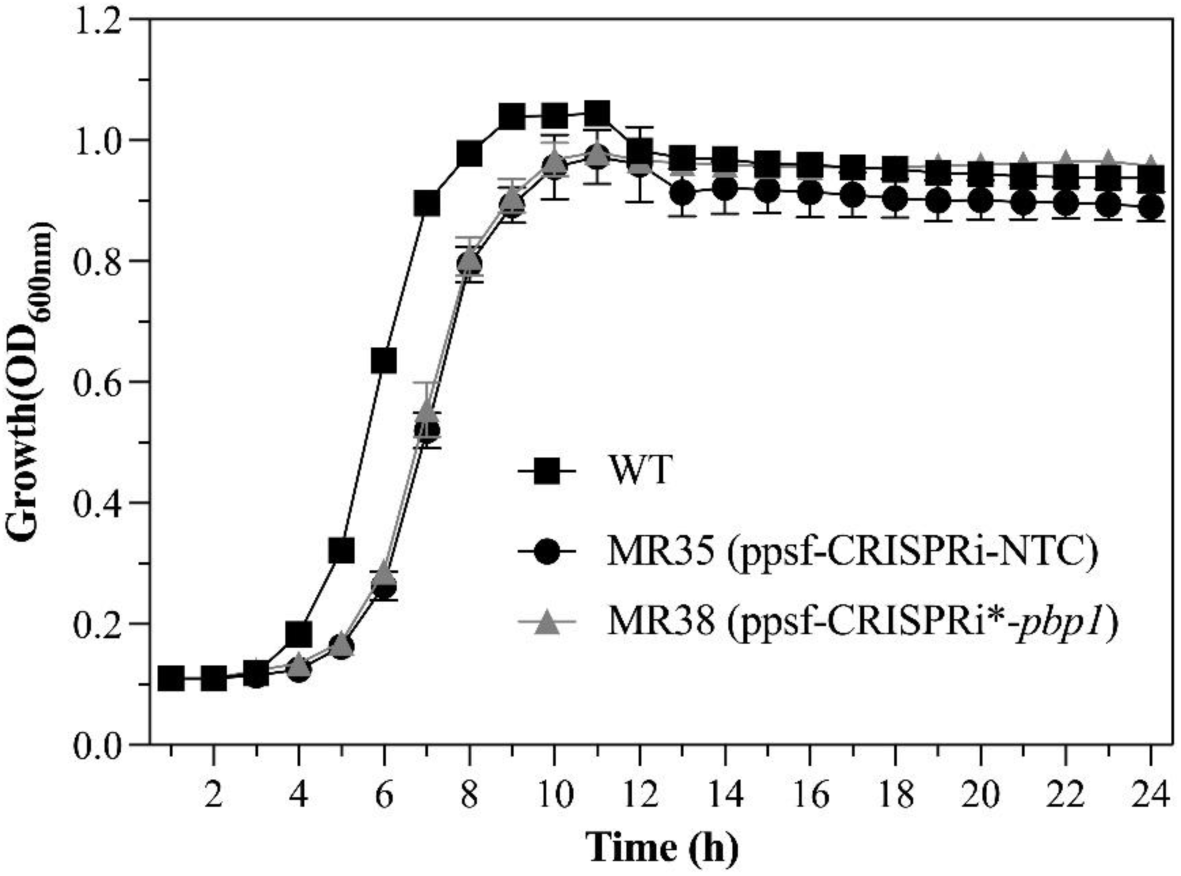
The ppsf-CRISPRi construct with a truncation in *dcas9* is not functional. Growth (OD600) of the indicated *S. aureus*, WT, ppsf-CRISPRi strains grown in TSB without chloramphenicol for WT and under chloramphenicol selection for ppsf-CRISPRi strains. Full-length, 4107 bp (in MR35) or truncated *dcas9*, 559 bp (in MR38) expression was controlled by the P*coa* promoter. The sgRNAs targeted either the essential peptidoglycan biosynthesis gene *pbp1* to induce growth inhibition, or an NTC sequence derived from the luciferase gene. Data show mean ± SD of n = 3 biological replicates (each recorded with three technical replicates).

**Figure S3.**
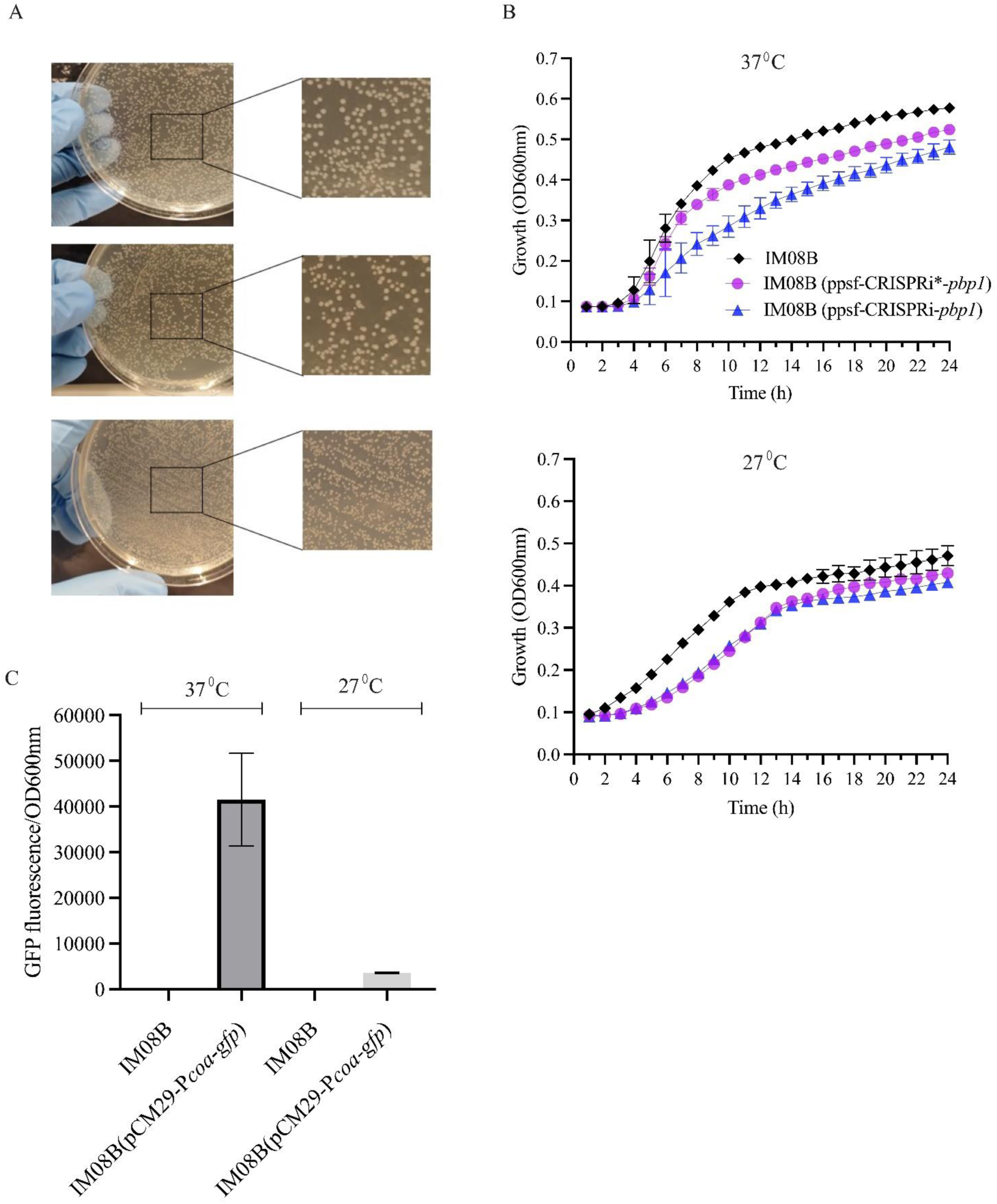
The toxicity of dCas9 is correlated with the level of dCas9 expression from ppsf- CRISPRi construct in *E. coli* IM08B. (A) Photographs of *E. coli* IM08B WT (top), IM08B transformants with ppsf-CRISPRi*(*pbp1*) (middle) and ppsf-CRISPRi (*pbp1*) (bottom) at 37°C. Small-colony phenotypes indicate cell toxicity (bottom). (B) Growth curves of *E. coli* IM08B (WT), IM08B transformants with ppsf-CRISPRi*(*pbp1*) and ppsf-CRISPRi (*pbp1*) at 37°C and 27°C. ppsf- CRISPRi* construct’s *dcas9* is truncated (559 bp) and nonfunctional. (C) *S. aureus* coagulase gene promoter (P*coa*) activity in *E. coli* IM08B at 37 and 27°C. *S. aureus* coagulase gene promoter-based fluorescent reporter plasmid (pCM29-P*coa-gfp*) was transformed into *E. coli* IM08B strains to check the level of functionality of that gene promoter by GFP fluorescence intensity measurement. GFP fluorescence data was normalized to the OD600 value of the corresponding sample. The *E. coli* IM08B strains without fluorescent reporter plasmid are included as controls. Bars show means ± standard deviation of n=3 biological replicates (each recorded with 3 technical replicates).

**Figure S4.**
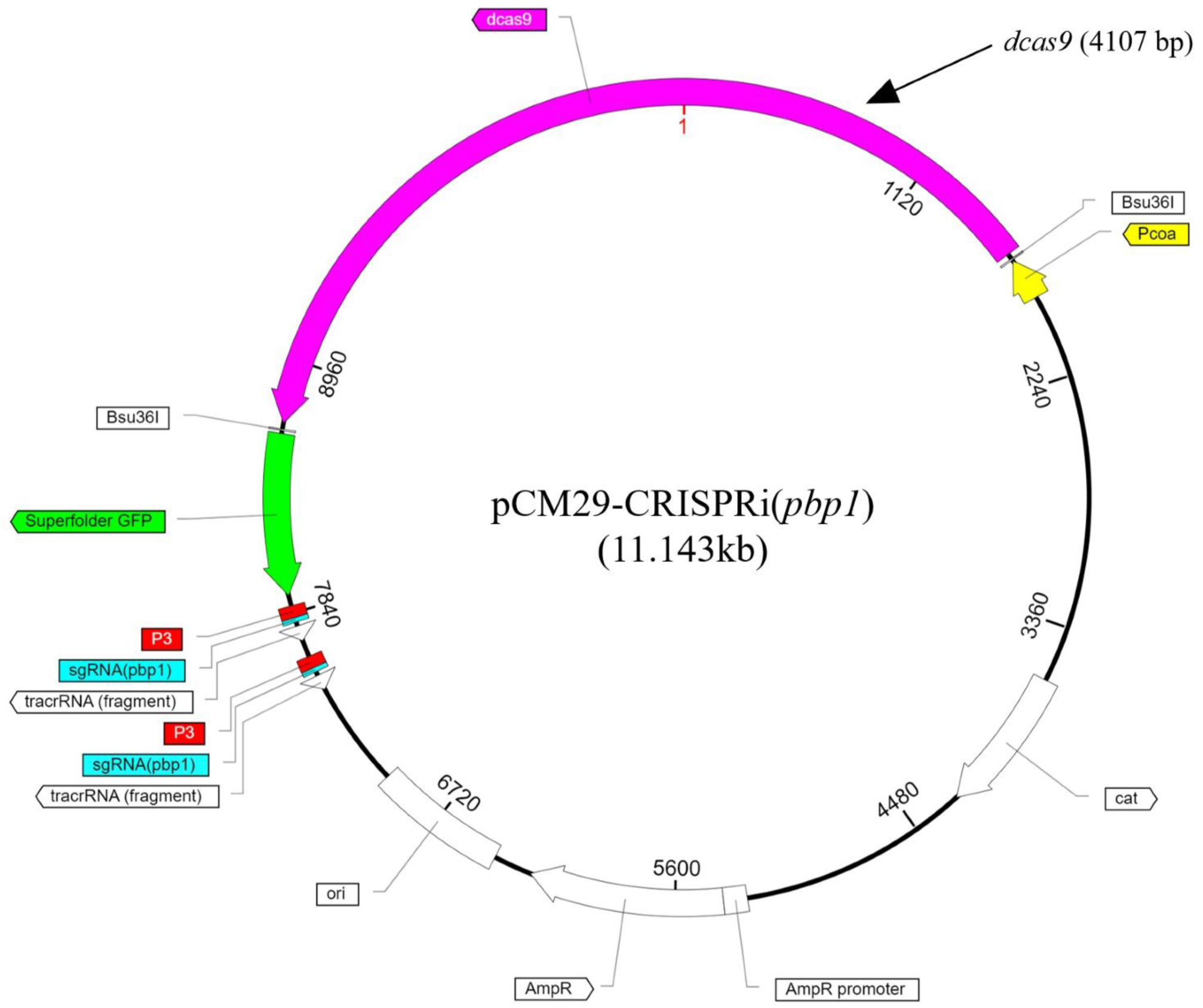
Stable ppsf-CRISPRi construct cloning in *E. coli* IM08B at 27°C. Plasmid map of ppsf- CRISPRi(*pbp1*), cloned in *E. coli* IM08B at 27°C. The plasmid was sequenced using nanopore techniques developed by Eurofins (https://eurofinsgenomics.eu/en/custom-dnasequencing/eurofins-services/whole-plasmid-sequencing/) and the plasmid map was created using SnapGene based on the provided whole plasmid sequence.

**Figure S5.**
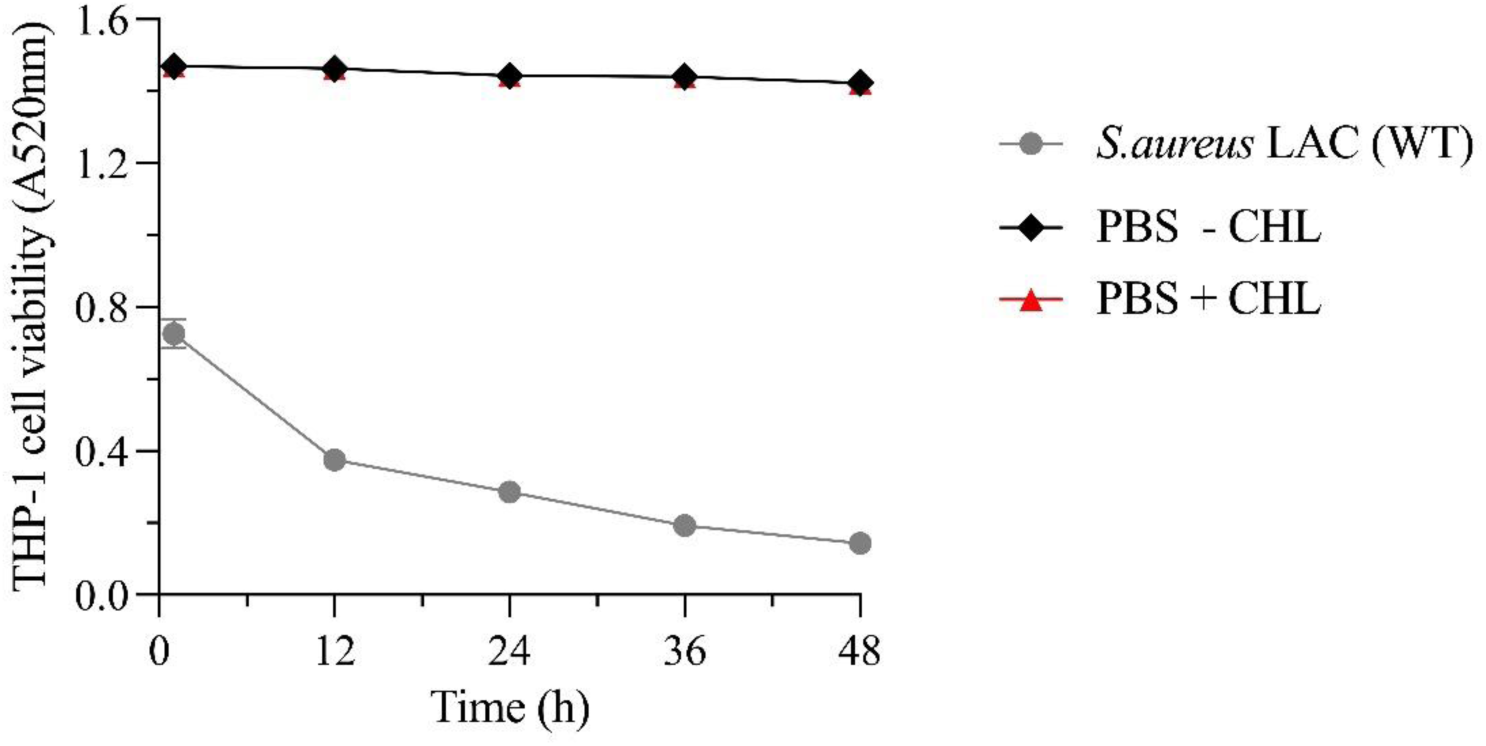
Chloramphenicol does not affect the viability of THP1 cells. Time-course THP-1 cell viability assay following infection with WT *S. aureus* USA 300 LAC strain or with PBS with (+) or without (-) CHL. THP-1 cells were incubated in RPMI+ with 50 µg/ml Gentamicin. The data represents the mean ± standard deviation of three biological replicates.

**Table S1.**
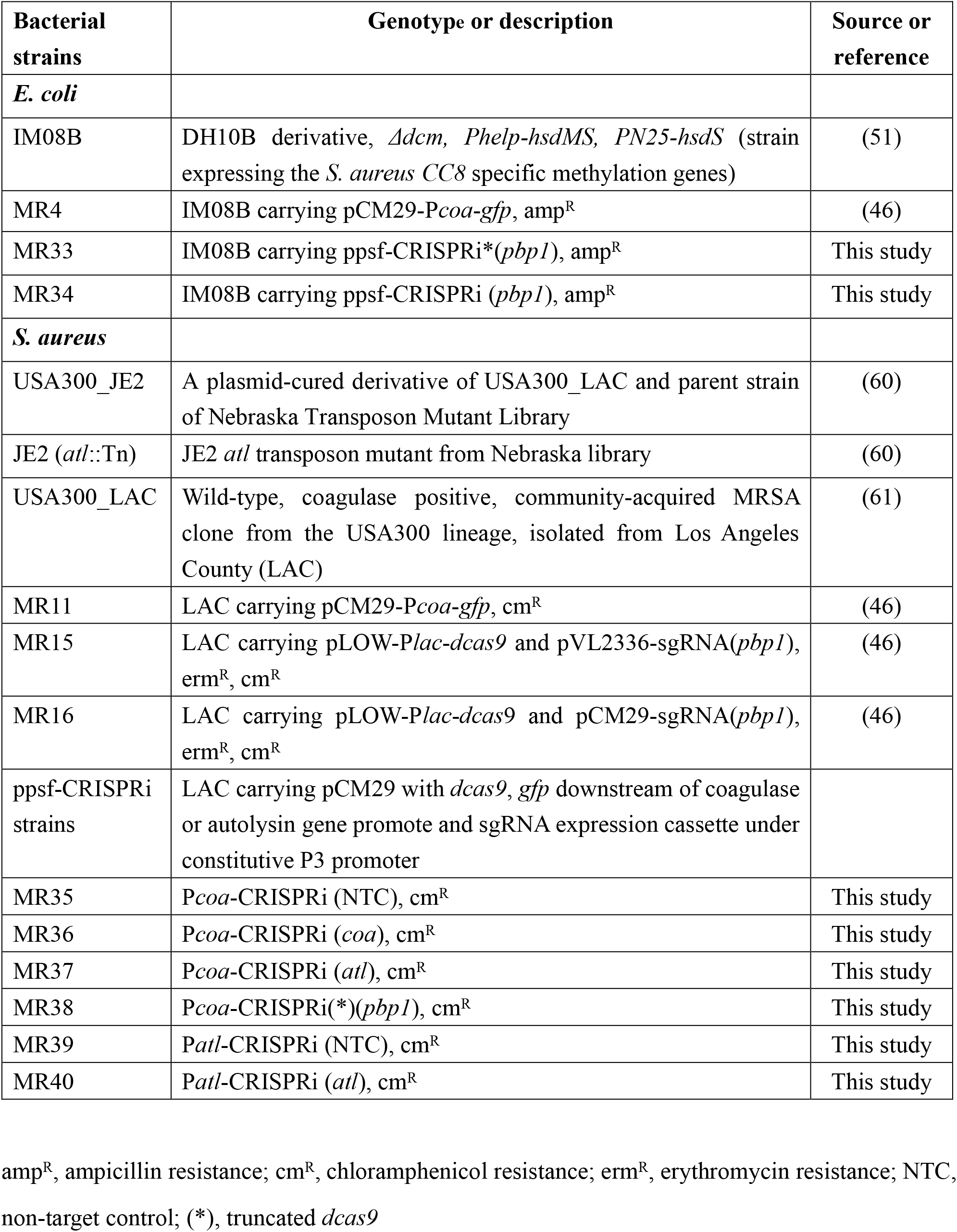
Bacterial strains used in this study.

**Table S2.** Plasmids used in this study.

| Plasmids | Relevant characteristics<sup>a</sup> | Source or reference |
| --- | --- | --- |
| pCM29-*Pcoa-gfp* | pCM29 carrying *gfp* downstream of *S. aureus* coagulase gene promoter<sup>b</sup>, amp<sup>R</sup>, cm<sup>R</sup> | (46) |
| pLOW- *Plac-dcas9* | pLOW carrying mutated Cas9 (*dcas9*) downstream of *lac* promoter, amp<sup>R</sup>, ery<sup>R</sup> | (18) |
| pVL2336-sgRNA (*pbp1*) | pVL2336 carrying sgRNA(*pbp1*) expression cassette, amp<sup>R</sup>, cm<sup>R</sup> | (18) |
| pCM29-sgRNA (*pbp1*) | pCM29 carrying sgRNA(*pbp1*) expression cassette, amp<sup>R</sup>, cm<sup>R</sup> | (46) |
| ppsf-CRISPRi constructs | pCM29 carrying *dcas9*, *gfp* downstream of *S. aureus* coagulase or autolysin gene promoter<sup>b</sup>, and sgRNA expression cassette under constitutive P3 promoter, amp<sup>R</sup>, cm<sup>R</sup> |  |
| *Pcoa*-CRISPRi(*) (*pbp1*) | sgRNA target *pbp1* | This study |
| *Pcoa*-CRISPRi (NTC) | sgRNA target NTC | This study |
| *Pcoa*-CRISPRi (*coa*) | sgRNA target *coa* | This study |
| *Pcoa*-CRISPRi (*atl*) | sgRNA target *atl* | This study |
| *Pcoa*-CRISPRi (*pbp1*) | sgRNA target *pbp1* | This study |
| *Patl*-CRISPRi (NTC) | sgRNA target NTC | This study |
| *Patl*-CRISPRi (*atl*) | sgRNA target *atl* | This study |

<sup>a</sup> amp<sup>R</sup>, ampicillin resistance; cm<sup>R</sup>, chloramphenicol resistance; ery<sup>R</sup>, erythromycin resistance; NTC, non-target control; (\*), truncated *dcas9*
<sup>b</sup> Promoter sequences were selected as the non-coding gap sequence between the gene and its upstream gene.

**Table S3.**
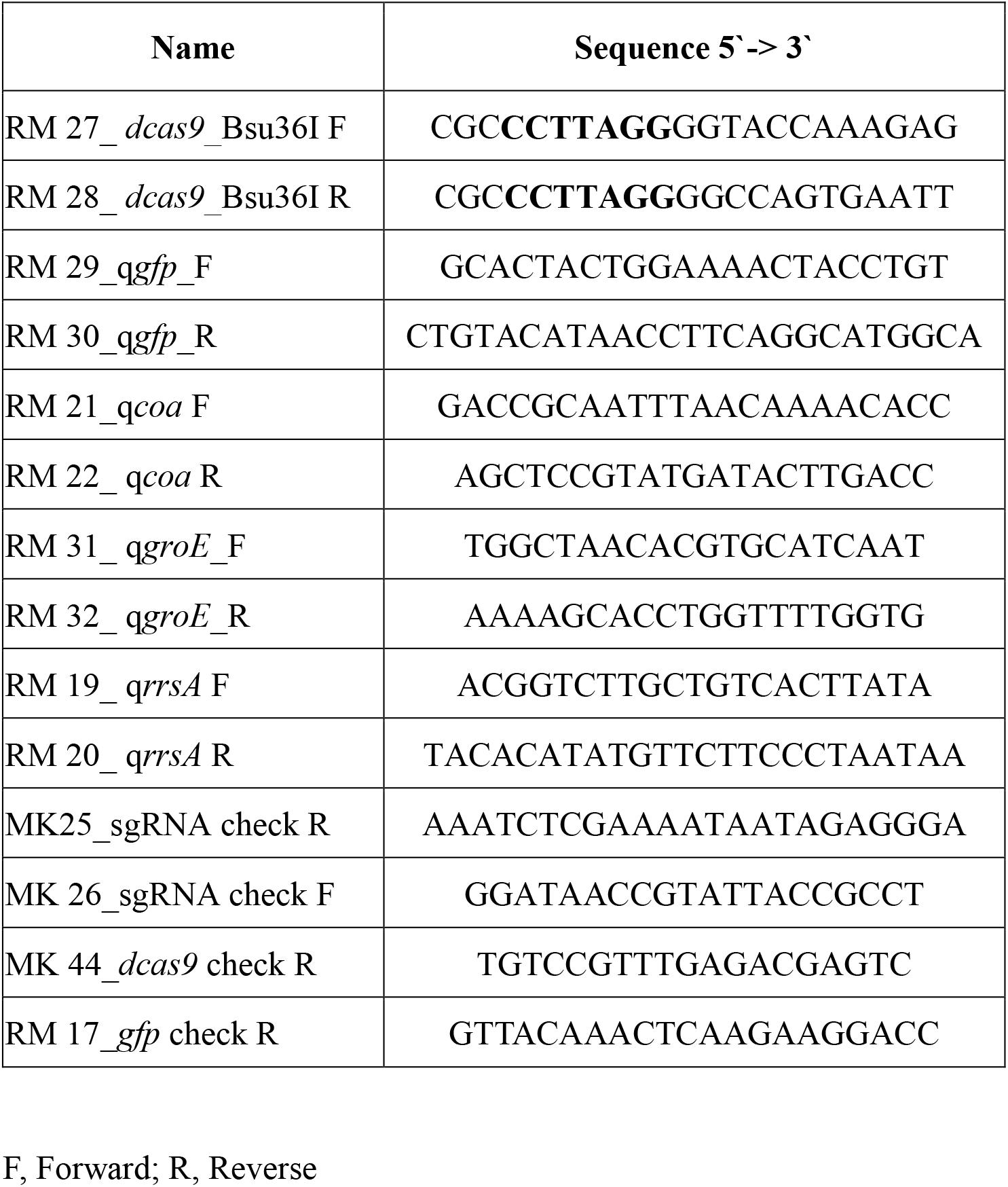
Primers used in this study.

**Table S4.**
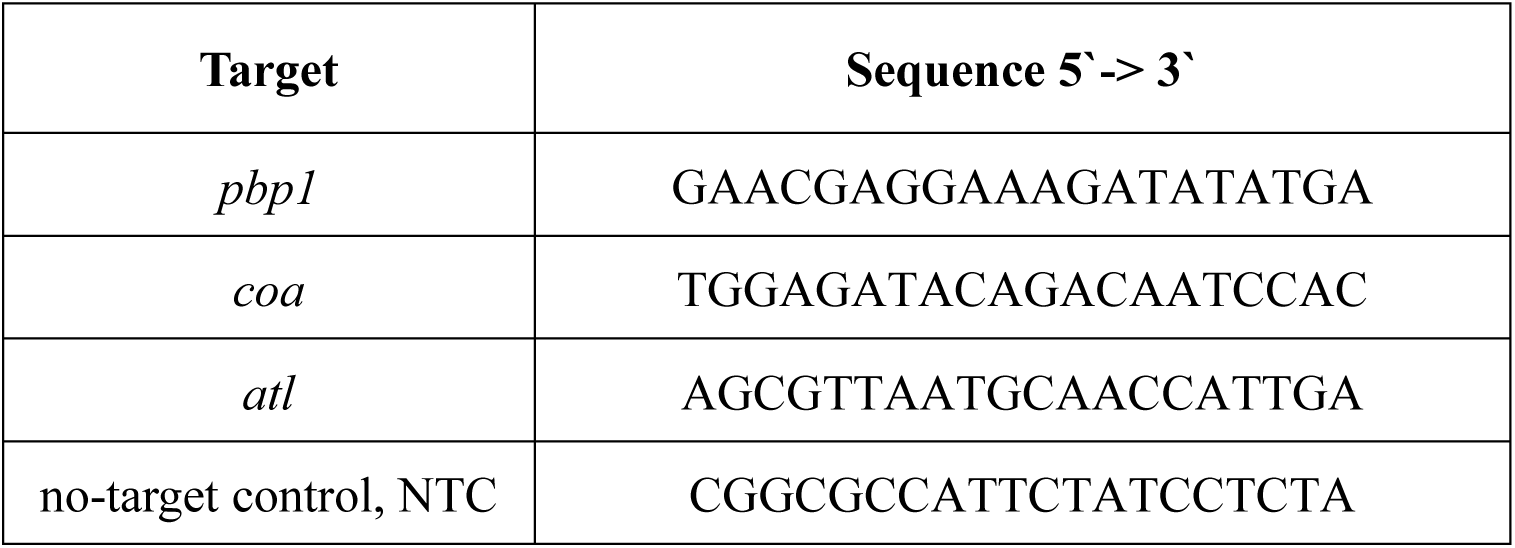
Sequences of sgRNA base pairing regions.

